# Information transfer in QT-RR dynamics: Application to QT-correction

**DOI:** 10.1101/269670

**Authors:** Ilya Potapov, Joonas Latukka, Jiyeong Kim, Perttu Luukko, Katriina Aalto-Setälä, Esa Räsänen

## Abstract

The relation between the electrical properties of the heart and the beating rate is essential for the heart functioning. This relation is central when calculating the “corrected QT interval” — an important measure of the risk of potentially lethal arrhythmias. We use the transfer entropy method from information theory to quantitatively study the mutual dynamics of the ventricular action potential duration (the QT interval) and the length of the beat-to-beat (RR) interval. We show that for healthy individuals there is a strong asymmetry in the information transfer: the information flow *from RR to QT* dominates over the opposite flow (from QT to RR), i.e. QT depends on RR to a larger extent than RR on QT. Moreover, the *history* of the intervals has a strong effect on the information transfer: at sufficiently long QT history length the information flow asymmetry inverts and the RR influence on QT dynamics weakens. Finally, we demonstrate that the widely used QT correction methods cannot properly capture the changes in the information flows between QT and RR. We conclude that our results obtained through a model-free informational perspective can be utilised to improve and test the QT correction schemes in clinics.

## Introduction

**The temporal coordination between the beating rate and the action potential propagation is vital for the proper function of the heart. Therefore, a quantitative measure of the relation between the action potential duration and the interbeat interval has potential applicability from both scientific and clinical points of view^1^.**

It is known that the QT interval of the electrocardiogram (ECG) recordings relates to the ventricular action potential duration and varies with the heart rate (HR)^1–4^. The relation between QT and HR has a practical importance as the assessment of the QT duration with respect to HR facilitates stratification of patients with the increased risk of severe arrhythmias, including different ventricular arrhythmias and *torsades de pointes*, which may result in sudden cardiac death^1, 4–7^. Additionally, the QT values normalised according to the HR variability are used in assessment of adverse cardiac side effects of drugs in the context of diseases not directly related to the heart, as cancer and others^1, 8, 9^.

Different formulas and methods to adjust the QT values according to the HR have been proposed. The most frequently used ones are the formulas of Bazett and Fridericia^10, 11^. Although the formulas result from simple fitting procedures, they are still widely used in medical practice^3, 12, 13^. The formulas have been criticised due to their inability to universally capture the QT-HR relationship^14, 15^, their subjective character^16, 17^ and extreme simplicity^17–19^. Some other approaches exist too, including graphical procedures^20^ and regression type corrections^1, 15^.

Additionally, recent evidence demonstrates improvements in the QT correction procedures when a range of the past values of HR is taken into account^4, 21, 22^. The correction improvement is especially profound in patients with atrial fibrillation, where HR values are unpredictable^23, 24^.

In this work, we analyse the coupled QT and beat-to-beat (RR) interval dynamics in terms of information flows. We use the transfer entropy method^25^ which quantifies information transfer between two related processes. Specifically, we demonstrate asymmetry in the information flows between QT and RR time series of the same individual. Moreover, we show that the history length of the past values significantly affects the information flows, and the effect is opposite when QT and RR history lengths are varied independently. Additionally, the RR-QT reciprocal configuration, where the QT interval may correspond to the RR interval of the preceding or the same beat, affects the information flow. Moreover, we demonstrate that the widely used QT correction formulas are not able to reduce the observed information transfers as stipulated by the correction procedures. Finally, we formulate an informational basis of the QT-correction for testing and developing of the QT-correction schemes.

## Methods

**Transfer entropy (TE) estimates information transfer (defined in Shannon terms) from the *source* process to the *destination* process. This is achieved by calculating the amount of information that the source preceding samples provide about the next sample of the destination in the context of the destination preceding samples^25^.** Figure 1 schematically shows the effects of the preceding values (history) on the next (*i*-th) value of a signal. For example, the RR→QT (green lines) information transfer is calculated by first taking only *k* QT history values and estimating their share in *predicting* the *i*-th QT value (dashed line) and, second, by considering both the share of *k* QT history values and the share of *n* RR history values in predicting the *i*-th QT value (solid line). Finally, the difference between the two estimations is directly related to the RR QT transfer^25, 26^.

**Figure 1.**
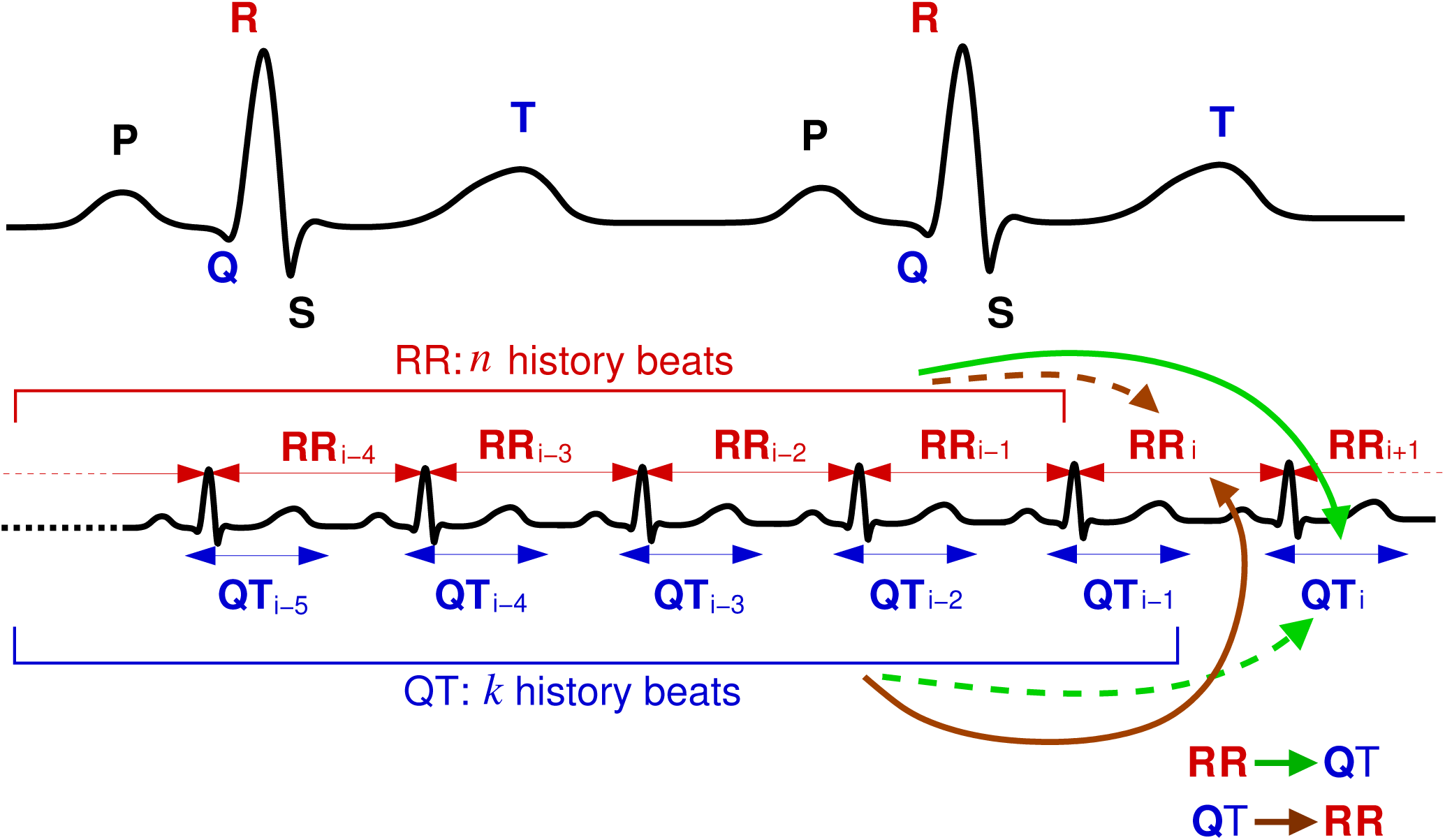
Information transfer between RR and QT series accounting for the preceding values (history). The two contributions are compared when calculating TE: the contribution of the *destination* history (**dashed** green/brown arrow) and that of the *source* history in the context of the destination history (**solid** green/brown arrow).

The relation between the preceding samples and the next sample of, e.g., QT series in the RR→QT transfer estimation is formally written as^25^:

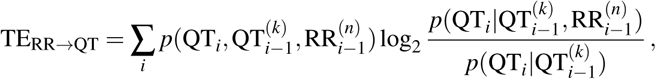

where *p*(*x*) is a probability distribution, *p*(*x|y*) is a conditional probability, 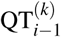 and 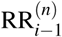 are *k* and *n* preceding values (from *i* – 1 backwards) of QT and RR series, respectively. The sum is taken over all states ^*i* 1^ of the process. The resulting TE measure is in bits of information and is the average over the process realisations. In this study we vary RR history (*n*) and QT history (*k*) lengths separately. Unless specified otherwise, the history length is one heartbeat, i.e. *n* = 1 and/or *k* = 1.

We use Java Information Dynamics Toolkit (JIDT version 1.3.1) for TE calculations^26^ with Kraskov-Stögbauer-Grassberger probability estimation^27^ (algorithm no. 1), 4 nearest neighbours, and default values for other parameters.

**We have tested some known synthetically generated time series with the TE method. The results of the tests are presented in the Supplementary Information, SI, Sec. S3.**

## Data

We downloaded raw ECG signals from PhysioNet^28^ (“MIT-BIH Normal Sinus Rhythm” and “MIT-BIH Long Term” databases). Then, we extracted RR and QT intervals using the provided software^28–31^ and discarded low quality signals. The final data set contains ECG recordings of 18 healthy individuals: 11 women (age from 20 to 45 years, mean 32), 7 men (age from 26 to 75 years, mean 46). Data were preprocessed to remove artefacts in the detected QT and RR values (see also Sec. S1, SI).

## Data Availability

All data analysed during the current study are publicly available at PhysioBank — an online archive of physiological signals maintained by PhysioNet. The data are available at: https://physionet.org/physiobank/database/nsrdb/ (“MIT-BIH Normal Sinus Rhythm”) and https://physionet.org/physiobank/database/ltdb/ (“MIT-BIH Long Term”).

## Results

### Information transfer asymmetry

First, we observe that the RR→ QT transfer is larger than the opposite QT→ RR transfer (Fig. 2). Since the TE algorithm adds a small amount of noise to the original data^26, 27^, we run TE calculations for 100 times, each time taking the average from 18 coupled time series in order to estimate the true mean value of the subject group. The resulting average TE distributions for the two information transfers are shown in Fig. 2A **(the standard deviation distributions are shown in Sec. S2, SI)**. The distributions are significantly (*P* < 0.0004 unpaired two-sided t-test) different. Moreover, the individual comparisons between 100 TE distributions with paired two-sided t-test result in the P-value distribution shown in Fig. 2B, which also confirms the significance.

**Figure 2.**
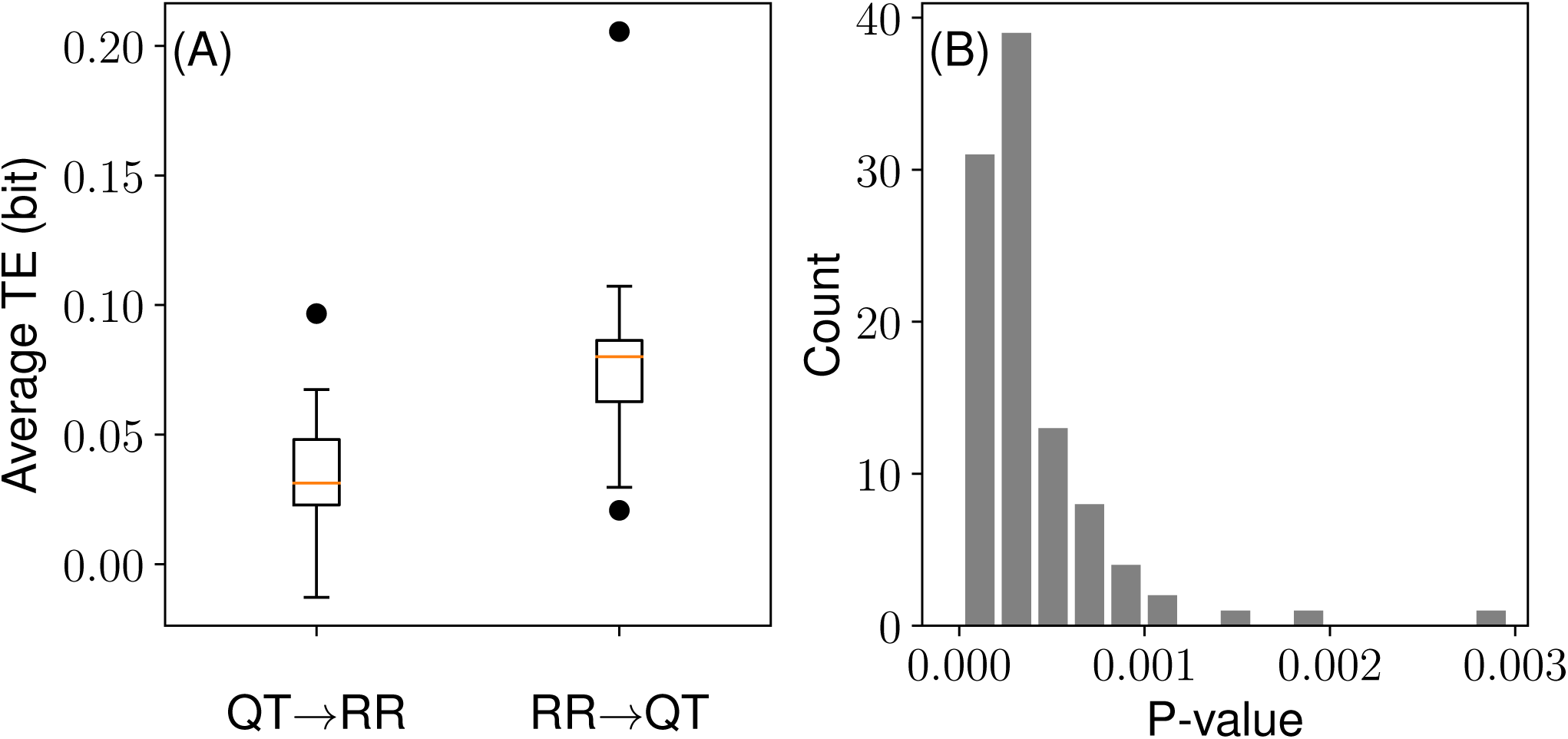
RR→ QT information transfer is larger than that of QT RR (*n* = *k* = 1). (A) Distribution of TE averages. The red lines show the median of the distributions. The upper and lower boundaries of the box depict the 1st and the 3rd quartiles of the distributions. The lines extend further to cover additional 1.5 of the interquartile range (length of the box). Circles (●) depict outliers (i.e. points beyond the covered ranges). (B) Individual comparison P-values (paired two-sided t-test) between the TE distributions.

The inequality TE_RR→QT_ > TE_QT→RR_ suggests that there is an implicit asymmetry in these otherwise equally important subprocesses in the context of the normal heart functioning. As TE_X→Y_ quantifies the degree of a dependency *Y* = *f* (*X*), the larger value of TE_RR→QT_, compared to TE_QT→RR_, indicates that the QT interval is more dependent on RR interval values, than RR on QT.

**Note that in this work by “influence” or “dependency” we do not mean causality, but rather correlation between the coupled processes. Thus, for example, a third process might be involved in causal effects between QT and RR (see Sec. S6, SI).**

#### Heartbeat history

Next, we consider effects of the history length of the RR/QT series on TE. For this, we vary the history length of each series (RR or QT) separately, while keeping the history length of the other signal minimal, i.e., one heartbeat.

Figure 3 demonstrates that with increasing history length the initial gap between RR→QT and QT→RR transfers increases further. However, as RR history length *n* becomes longer, TE_RR→QT_ starts dominating over the opposite transfer; while as QT history length *k* increases, TE_QT→RR_ becomes larger than the opposite transfer. Note that TE_RR→QT_ at large *n* is still larger than TE_QT→RR_ at large *k* (Fig. 3). Additionally, TE_QT→RR_(*n*) reaches zero after 20…25 beats, while TE_RR→QT_(*k*) is not effectively zero at the farthest point considered (*k* = 50). These results indicate that the RR time series have a strong influence on the QT series as well as a larger informational content.

**Figure 3.**
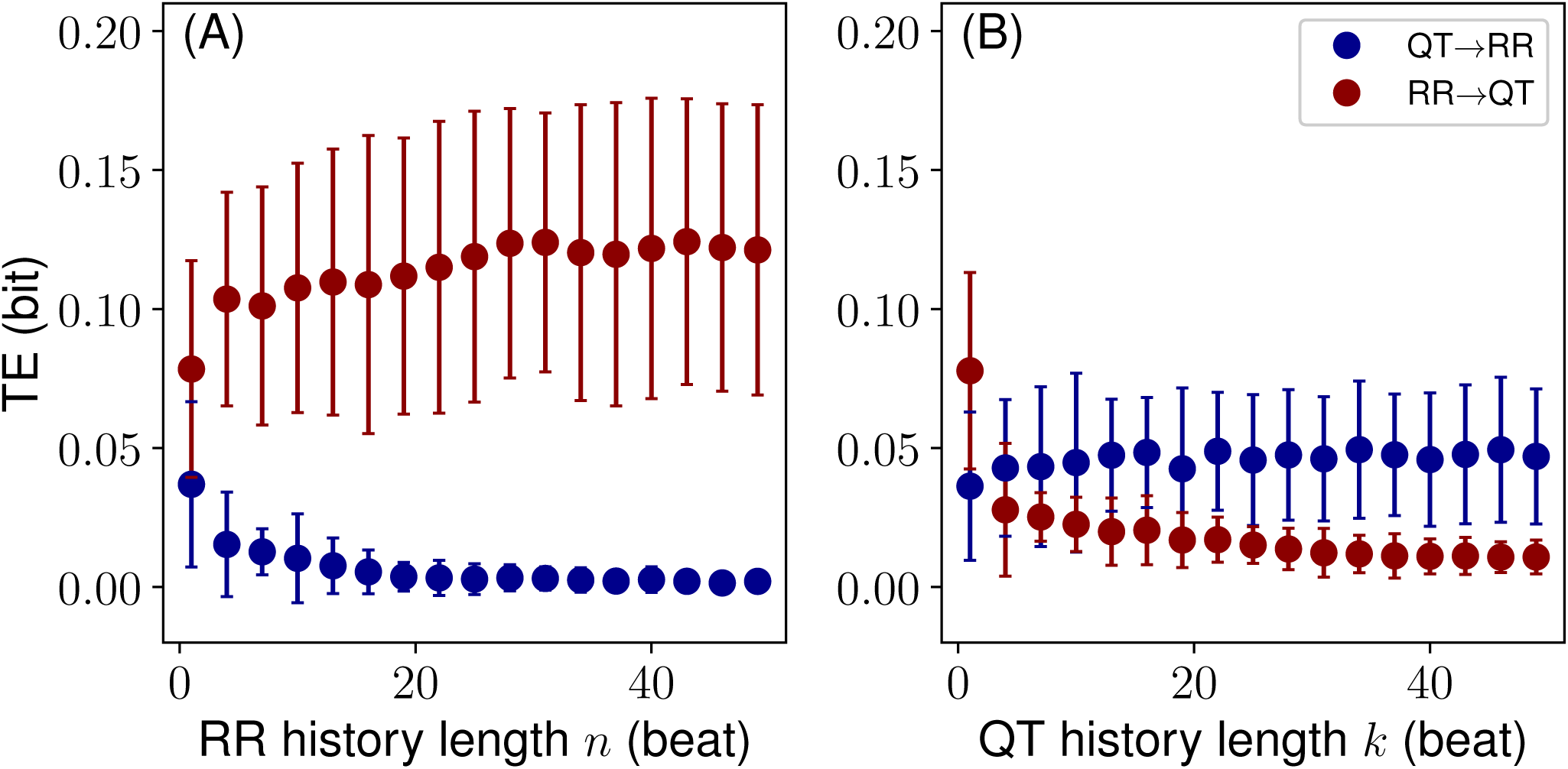
History length effect on RR→QT and QT→RR information transfers: (A) the RR history *n* is varied (*k* = 1); (B) the QT history *k* is varied (*n* = 1). Dots (●) represent mean values, whereas the error bars standard deviations.

Finally, in Fig. 3 we can also determine the *critical* history length, *n*_*crit*_, after which the information transfers remain constant. Thus, for example, TE_RR→QT_ reaches a plateau at around 20…30 beats of RR history. This critical value reveals a characteristic length of the RR history which is able to explain the QT variability. **Importantly, the critical length does not directly relate to the real heartbeat history because during the preprocessing different segments of consecutive beat ECG‘s were concatenated to form a single time series per individual. Nevertheless, this result demonstrates that the characteristic history length exists. Moreover, for the consecutive heartbeat segments** *n*_*crit*_ **can be seen as well (see Sec. S1.3, SI).**

In order to verify that the history dependence in fact takes place as shown in Fig. 3, we conduct a permutation test, where QT and RR series from different individuals are mixed to form new unrelated coupled time series. Here, we use 10 permutations, resulting in 180 coupled QT/RR series. We calculate the TE distributions over different RR and QT history lengths and then compare them to the original distributions corresponding to Fig. 3. The comparison in the form of P-values (unpaired t-test) is shown in Fig. 4. The results confirm that all transfers arise from the genuine QT/RR relationship in the individuals, that is, the original transfers are significantly different from the surrogate permuted QT and RR sequences. The QT→RR transfer might look as an exception to the rule; however, TE_QT→RR_ becomes apparently zero after several beats of the RR history *n* (Fig. 3), indicating that there is no information transfer.

**Figure 4.**
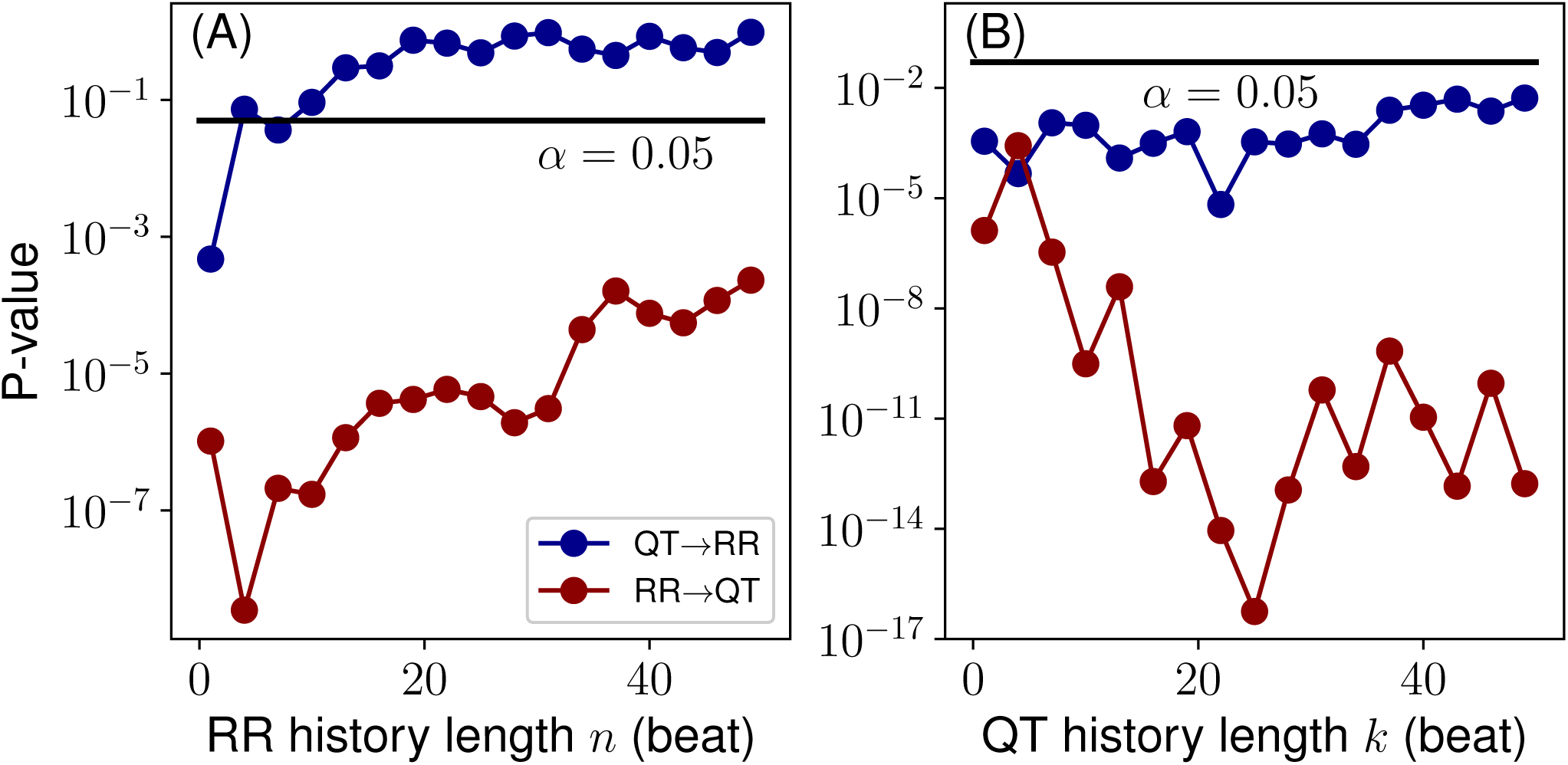
P-values resulting from (unpaired) t-test comparisons between 18 true and 180 permutation distributions for each history length value (cf. Fig. 3): (A) for varying RR history, *n* (*k* = 1); (B) for varying QT history, *k* (*n* = 1). Solid black line denotes the false discovery rate *α* = 0.05.

#### QT-RR coupling configurations

A coupled QT/RR process can assume two configurations in regards to how the QT interval is selected. The QT interval may be coupled with **the *preceding* RR interval (Fig. 1). This configuration was used in all results reported in this work and seems to be the most common choice of the heart research community (see, e.g., Ref.^32^). In the alternative configuration, however, the QT interval is coupled with the RR interval of the same heartbeat (Fig. 5).** In this section, we study whether the configuration can cause differences in the information transfers between RR and QT (Fig. 6).

**Figure 5.**
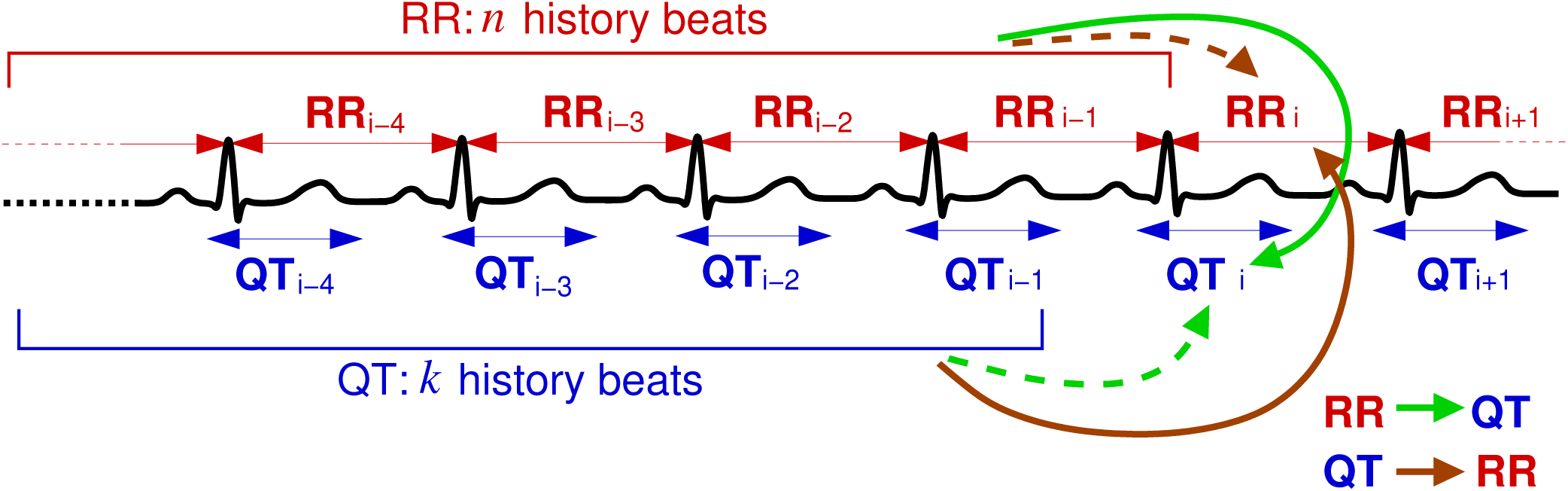
A schematic representation of the coupling configuration, where QT is coupled with the same heartbeat RR. Compare with the configuration, where QT is coupled with the preceding RR (Fig. 1).

**Figure 6.**
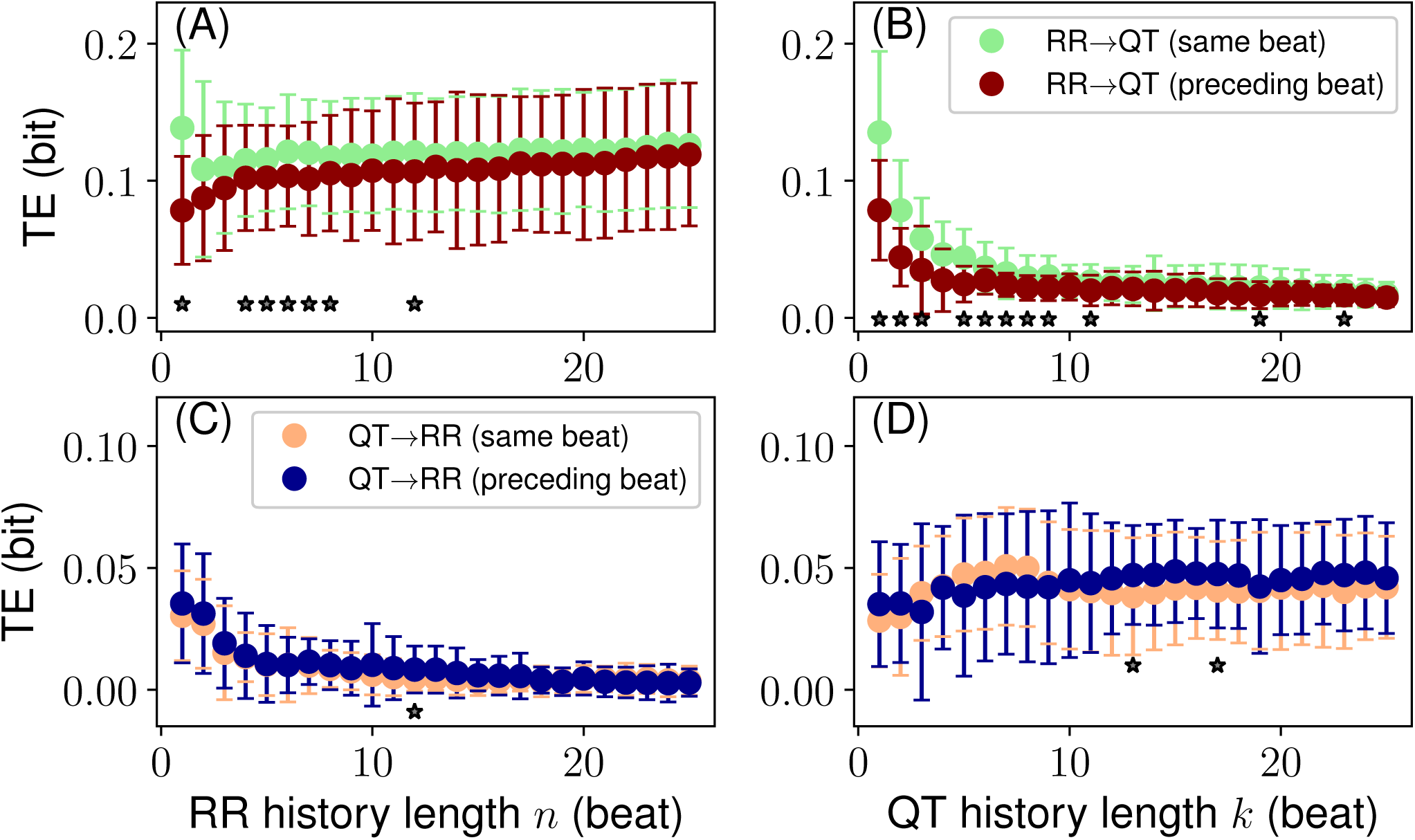
Information transfers for two coupling configurations: QT with the preceding heartbeat RR (“preceding beat”) vs. QT with the same heartbeat RR (“same beat”). (A) and (B) TE_RR→QT_ over *n* and *k*, respectively. (C) and (D) TE_QT→RR_ over *n* and *k*, respectively. Stars show significant differences between distributions (*P* 0.05, paired t-test). Format: mean standard deviation.

On the one hand, TE_RR→QT_ significantly differs between the two configurations at small *n* and *k* with the mean transfer for the “same beat” configuration being larger than the mean transfer for the “preceding beat” configuration. The effect is especially pronounced for several first beats of the history (Fig. 6). On the other hand, TE_QT→RR_ does not reveal such tendencies, although occasionally TE distributions at some history lengths seem to be significantly different (Fig. 6).

**The configuration with the same heartbeat coupling reveals larger information transfer than the preceding heart-beat coupling due to the effective shortening of the history by one heartbeat in the case of the former. Thus, the latter is weaker for** TE_RR→QT_ **as the history influence vanishes with the length. Overall, the preceding-beat configuration introduces the delay of one heartbeat to all history lengths. This is especially evident in the case of one beat history, where** TE_RR→QT_ **for the same beat configuration is significantly larger than the similar transfer for the opposite configuration (Fig. 6). This finding suggests the strong effect of the history on the information transfers as well as its non-uniformity over the history length. Moreover, it affirms the importance of RR of the first preceding beat in determining QT (this also has been found on a class of exponentially weighted models in Ref.^22^.**

The RR→ QT transfer, being a direct measure of QT dependence on RR, is affected by the mutual QT/RR configuration. The QT correction schemes must account for this, especially if the procedure is based on the previous beats as in Refs.^4, 21^ and given that the effect is not uniform along the history length.

#### Gender effects

It is known that women have longer QT than men^33–35^. We have selected male and female subgroups from the studied cohort and calculated the TE distributions for different history lengths. Although we did not find any significantly persistent discrepancies between the transfers, the TE_RR→QT_ distributions over *n* demonstrate systematic deviations. Namely, the TE_RR→QT_ mean value is persistently higher for males than for females at all *n*. We emphasise that our findings are based on a small group of 7 men and 11 women. Therefore, further studies with a larger group size might be needed (see also Sec. S5, SI).

### Informational basis of the QT-correction

The QT interval correction is a known technique for normalising the QT interval duration for different patients in order to be able to compare them at different heart rates and to determine whether there are signs for the increased risk of potentially lethal arrhythmias^1^.

Here, we correct the observed QT time series with several frequently used QT-correction procedures^10, 11, 22^ and calculate the TE distributions at various history lengths for the corrected QT, QTc, and RR time series.

In Fig. 7 we show the results obtained using the Fridericia formula. Other procedures including the use of the Bazett formula, RR history averaging, and exponentially weighted average of the RR history are presented in SI (Sec. S4).

**Figure 7.**
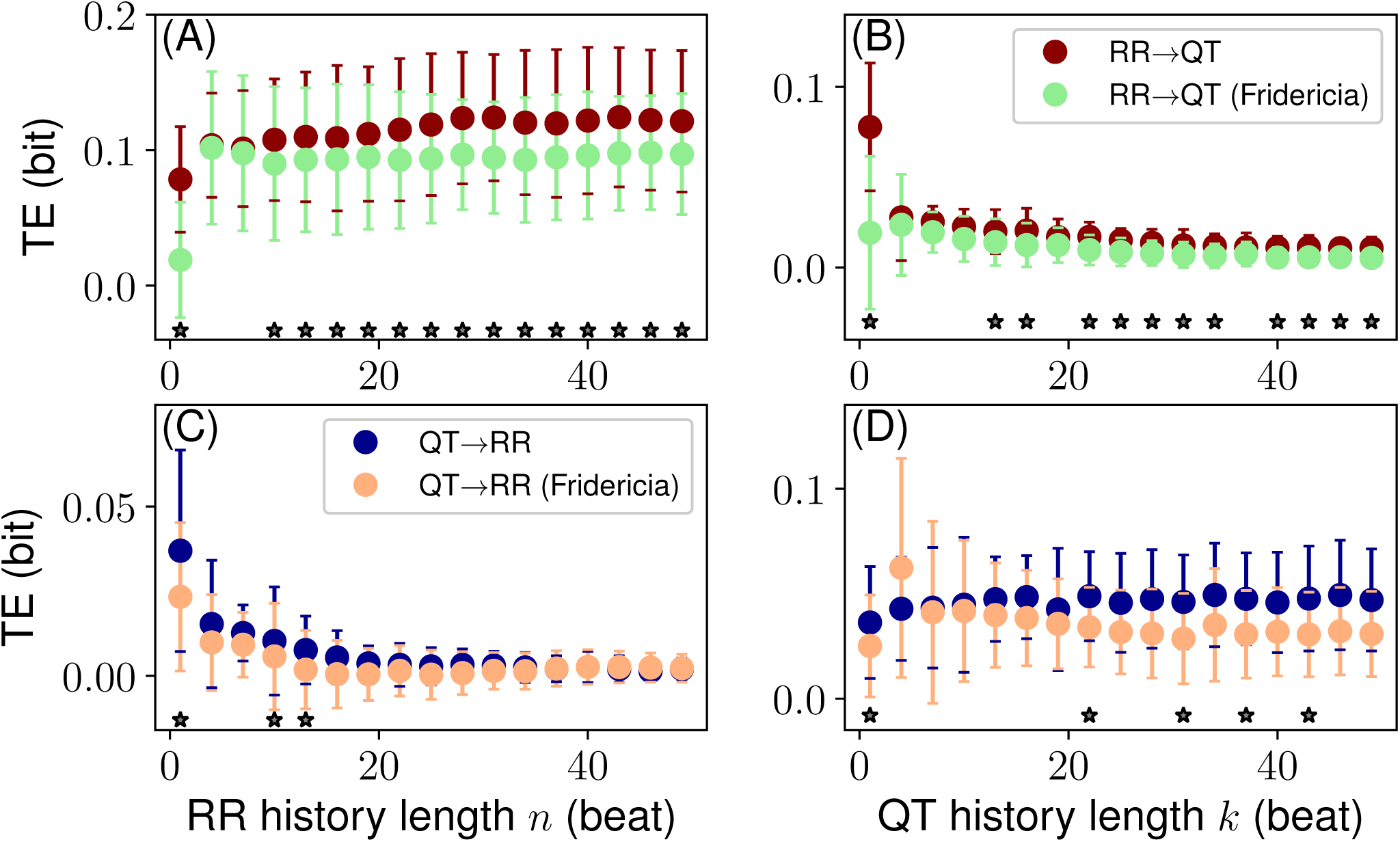
The effect of the QT correction (Fridericia formula) on the information flows. (A) and (B) TE_RR→QT_ over *n* and *k*, respectively. (C) and (D) TE_QT→RR_ over *n* and *k*, respectively. The stars show significant (*P ≤* 0.05, paired t-test) differences between TE distributions of the original and corrected signals. Data in the format mean *±* standard deviation.

For the Fridericia correction we find that all transfers are significantly different at *n, k* = 1 (the same result holds for Bazett correction, Sec. S4.1, SI). The corrected RR→ QT transfer deviates systematically from the original transfer at *n, k >* 10 *…* 12 beats towards zero (the systematic deviations were not observed for Bazett correction, Sec. S4.1, SI). The corrected QT→RR transfer is statistically similar to the original transfer at most of the history lengths, both *n* and *k* (Fig. 7).

We note that the purpose of the QT correction procedures is to eliminate the dependence of QT on RR. Thus, the corrected values of QT should ideally show no inter-dependency with RR. Hence, information transfers must be equal to zero. However, in Fig. 7 we observe significant non-zero transfers for the QT corrected series, though the tendency towards zero is evident in the case of the Fridericia-corrected RR→ QT transfer as the history length increases. Such a tendency is absent in the case of the Bazett correction, where all transfers are almost always statistically similar, except at *n, k* = 1 (Sec. S4.1, SI).

Finally we have tried two other QT-correction strategies, in which the Fridericia and Bazett formulas are used too, but the effective RR value in the formulas is calculated by means of: i) the average of the history of RR, and ii) the exponentially weighted average of the history of RR. The results are reported in Secs. S4.2 and S4.3, SI.

These results, being still not satisfactory in terms of the correction requirements to the RR→QT transfer, show interesting traits. Namely, the Fridericia correction reduces the TE_RR→QTc_ more than the Bazett correction. Moreover, TE_RR→QTc_ has a pronounced minimum around *n* = 20 heartbeats for the RR history averaging model (Sec. 4.2, SI). Additionally, the exponentially weighted average model shows significant reduction in RR→QTc (Fridericia formula) transfers with minimum occurring around *n* = 20 heartbeats. The best reduction is achieved when less contribution of the immediately preceding beat (larger contribution of the exponentially weighted history) is considered and for larger time constants (only for the Bazett formula), determining the rate of exponential decay of the weights (Sec. 4.3, SI). Similarly, the best reduction occurs at around *n* = 20 heartbeats. Both these results are in agreement with the *critical* history length, that is 20 beats of the RR history is an internal parameter of the coupled QT/RR time series used in this work.

#### Dynamical QT-correction

In order to eliminate the QT dependency on RR one needs to attain the effective absence of the information flow from RR to QT, that is TE_RR→QT_ = 0. So far we have considered the average TE over the time series. However, the local TE values form a time series where each point corresponds to a point in the QT/RR coupled time series. Decoupling of the QT and RR time series for each time point would mean finding, given the history, a QT/RR pair that leads to TE_RR→QT_ = 0 (Fig. 8A).

**Figure 8.**
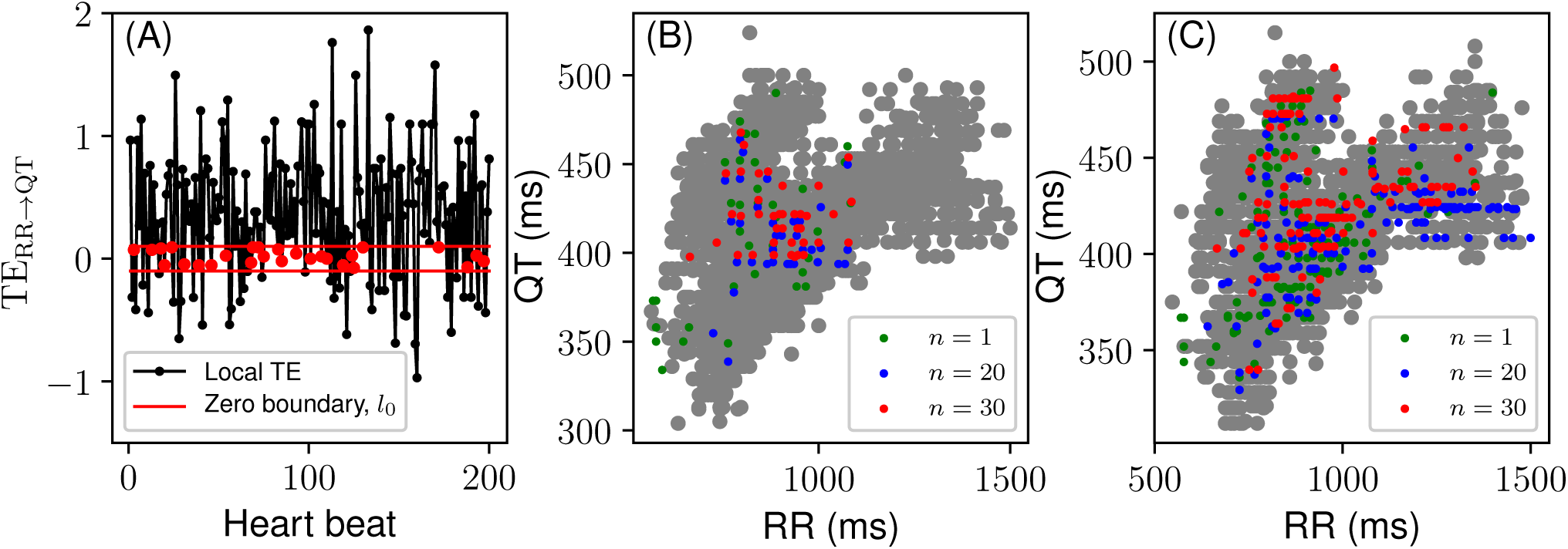
Dynamical QT-correction. (A) Local transfer entropy with values falling within the zero boundary, *l*_0_ (in red, for demonstration purposes *l*_0_ is larger than the actual value used in calculations). (B) QT/RR points corresponding to |TE_RR→QT_| < *l*_0_ for different RR history length, *n*; *k* = 1; *l*_0_ = 0.0001 bit (little noise is added to the position of the points for better resolution). (C) Same as (B) but *l*_0_ = 0.001 bit. Grey points show QT/RR values corresponding to |TE_RR→QT_| *>* 0.1 bit. The result is cumulative from all individual QT/RR time series.

Now, we can formulate a transfer entropy based QT-correction procedure:

*At every time sample in the TE (RR→ QT) series find a QT value for that sample so that the TE value becomes (effectively) zero. The found QT value is the QT-corrected interval.*

Note that in this procedure one should change the QT interval only in order to preserve the correlation structure in the context of the time series. For example, the RR interval and/or history cannot be changed.

The proper TE-based QT-correction implementation requires further technical studies of the probability distribution estimation algorithm and its parameters (e.g. k-nearest neighbors) as well as the history effects in different contexts of healthy and pathological QT/RR time series. As these would go beyond this initial study, we have shown the scattering of QT/RR points with the corresponding zero RR→QT transfer. Since the whole group contains a small portion of such QT/RR pairs we found the points where |TE_RR→QT_| < *l*_0_, where the zero boundary *l*_0_ is a small constant. The results are presented in Fig. 8.

One can see that for *l*_0_ = 0.0001 bit the cloud of QT/RR points with TE_RR→QT_ *< l*_0_ gets more aligned along the RR-axis and less resembling the high TE cloud (grey) as *n* increases and approaches *n*_*crit*_ (Fig. 8B). Additionally, for *n* = 30 the cloud (red) stays localized within physiological limits for QTc. For *l*_0_ = 0.001 bit the clouds of zero-TE QT/RR points widen over both RR- and QT-axes and get closer in shape to the grey cloud (Fig. 8).

These observations need to be further confirmed with other tests to determine, for example, the contribution of TE_QT→RR_ to the QT dependency on RR, the precision of the probability estimation algorithm, and the sensitivity of TE_RR→QT_ (the value of *l*_0_).

## Discussion

In this work we have considered the transfer of information between coupled RR and QT interval dynamics of healthy individuals. We used the transfer entropy (TE) method for this purpose^25^. An important application of the analysis is the QT correction, which is a standard tool for diagnostic purposes of potentially lethal arrhythmias^1, 2^. To the best of our knowledge, this study is the first attempt to establish a firm information theory approach to this clinical problem.

We have shown that the information transfer from RR to QT is larger than the transfer from QT to RR in the simplest case of accounting only for the effect of one previous beat of history. Thus, there is a hidden asymmetry of the information flow between RR and QT time series. Moreover, the inequality indicates that the QT interval is more dependent on RR interval values than RR on QT. The former is implicitly entailed by numerous studies on the QT interval correction; see, e.g., Refs.^1–4, 10, 11^ and references therein.

The aforementioned asymmetry of the information flow becomes more profound as more RR past values are taken into account. This finding is in agreement with the recent studies suggesting that the longer RR history can improve the QT correction^21, 22^. Namely, the increase in the RR→ QT information flow with the RR history length indicates a stronger ability of the RR history to explain the variability in QT, hence stronger QT dependence on RR. However, if longer QT history is considered the information flows reverse, indicating the strong RR dependence on QT. This result emphasises the importance of comprehensive analysis of the mutual QT/RR dynamics, considering, for example, the QT history as well. However, this is usually neglected in the QT correction studies.

There is a characteristic history length *n*_*crit*_, such that for history lengths longer than *n*_*crit*_ the QT correction based on the RR history should not produce any improved outcome. That is, the RR intervals of the past beyond *n*_*crit*_ are no more able to explain the variability in QT since the RR→ QT transfer reaches the steady state. **Using our data, however, it is impossible to relate** *n*_*crit*_ **to the real history length. Nevertheless, this and the study on the consecutive heartbeat ECG segments (see Sec. S1.3, SI) reveal the** TE_RR→QT_ **steady state well below the 50 heartbeat limit. Thus, the effective history length used in many clinical practices (e.g. cardiac safety assessment of drugs) may differ from the current protocols when analysed with transfer entropy.**

In the literature, other attempts have been made to quantify the dynamic QT-RR coupling^4, 22, 36–38^. These approaches rely on statistical models of the given form and, thus, have assumptions behind the models. For example, in Ref.^4^ the authors proposed to use the weighted average of the preceding RR intervals to account for the RR history effect on QT. They found that on average 150 preceding beats are enough to model QT accurately. Unlike these approaches transfer entropy accounts for the history *unconditionally* **(i.e. there are no assumptions, regarding the history, for example, implied by averaging or filtering of any sort).**

**We have shown that the mutual coupling configuration between QT and RR affects the information transfer.** Since this transfer reflects the degree of dependency of QT on RR, one would expect that the choice of the mutual configuration will quantitatively affect the relevant QT correction procedures. However, so far this distinction was not brought to attention in the studies pertaining to the subject of the QT correction (the usual choice of the research community is the preceding-beat configuration, see, e.g., Ref.^32^).

We have not found any statistically significant differences between the male and female subgroups, although for our group the mean RR→QT transfer was systematically larger for males than for females.

Additionally, we have estimated the effects of the QT correction on the information transfer between QT and RR. For this, the widely used QT correction formulas (Fridericia^11^ and Bazett^10^) were analysed. We conclude that the most commonly used QT correction formulas cannot provide for the proper informational reduction in the corrected QT/RR time series for all lengths of the heartbeat history. **Finally, we have outlined a TE-based QT-correction scheme. Although there is no straightforward implementation readily available for the scheme, it can be used in *devising, improving*, and *testing* QT-correction methods.**

In conclusion, this purely empirical, theory-free, and probabilistic approach to the QT correction problem might prove to be more effective in the medical context than the conventional “best-fit” formulae, as it establishes quantitative measures, uses probabilistic features, and does not have any constraining presumptions of the model itself.

## Acknowledgements

We are specially thankful for useful discussions to Joseph Lizier and Jason Lloyd-Price. This work was supported by Academy of Finland, Key Project “Health tracking through fractal analysis of complex signals” [project number 304458].

## Author contributions statement

I.P. conceived the study, conducted all calculations, and wrote the manuscript, J.L. preprocessed the data, J.K., P.L. and K.A-S. analysed the results, E.R. conceived the study, analysed the results, and wrote the manuscript. All authors reviewed the manuscript.

## Additional information

The authors declare no competing interests.

